# Investigating the temporal dynamics and modelling of mid-level feature representations in humans

**DOI:** 10.1101/2025.03.18.643889

**Authors:** Agnessa Karapetian, Alexander Lenders, Vanshika Bawa, Martin Pflaum, Raphael Leuner, Gemma Roig, Kshitij Dwivedi, Radoslaw M. Cichy

**Affiliations:** Department of Education and Psychology, Freie Universität Berlin, Berlin, Germany; Charité – Universitätsmedizin Berlin, Einstein Center for Neurosciences Berlin, 10117, Berlin, Germany; Bernstein Centre for Computational Neuroscience Berlin, Berlin, Germany; Faculty of Biology, Albert-Ludwigs-Universität Freiburg, Freiburg, Germany; Fraunhofer Institute for Laser Technology ILT, Aachen, Germany; Department of Mathematics and Computer Science, Freie Universität Berlin, Berlin, Germany; Department of Computer Science, Goethe Universität Frankfurt, Frankfurt am Main, Germany; Berlin School of Mind and Brain, Faculty of Philosophy, Humboldt-Universität zu Berlin, Berlin, Germany

**Keywords:** Mid-level features, scene perception, encoding, EEG, CNN

## Abstract

Scene perception is a key function of biological visual systems. According to the hierarchical processing view, scene perception in the human brain begins with low-level features, progresses to mid-level features, and ends with high-level features. While low- and high-level feature processing is well-studied, research on mid-level features remains limited. Here, we addressed this gap by investigating when mid-level features are processed in humans using a novel stimulus set of naturalistic scenes as images and videos, accompanied with ground-truth annotations for five mid-level features (reflectance, lighting, world normals, scene depth and skeleton position), and two framing features: one low-level (edges) and one high-level feature (action). To reveal when low-, mid- and high-level features are represented in the brain, we collected electroencephalography (EEG) data from human participants during stimulus presentation and trained encoding models to predict EEG data from ground-truth annotations. We revealed that mid-level features were best represented between ∼100 and ∼250 ms post-stimulus, between low- and high-level features. Moreover, we assessed scene- and action-trained convolutional neural networks (CNNs) as models of mid-level feature processing in humans. We found a comparable processing order for mid-but not low- or high-level features with humans. Overall, our results characterize mid-level feature processing in humans in the temporal domain and reveal CNNs as suitable models of the processing hierarchy of mid-level vision in humans.

## 1. Introduction

Visual scene perception is a crucial function of biological systems which enables humans to navigate and make decisions in their environment. According to classical theories of scene perception, it takes place in a hierarchical manner (Epstein & Baker, 2019; Groen et al., 2013, 2017; Malcolm et al., 2016; J. M. Henderson & Hollingworth, 1999), starting with the extraction of low-level features such as edges, contrast and color at ∼50 ms post-stimulus (Di Russo et al., 2002), and culminating with high-level semantic features such as scene size at ∼250 ms (Cichy et al., 2017) and action identity at ∼400 ms (Ge et al., 2019).

While the processing of low- and high-level features in the brain is relatively well-understood, there remains a gap in the understanding of the intermediate transformations, i.e., of the mid-level features. These are features related to our perception of surfaces and shapes in our environment (Anderson, 2020), involved in the construction of 3D representations that form the basis for high-level vision tasks (Biederman, 1972; Marr, 1982). Previous research focused on outline-related mid-level features such as texture (Long et al., 2018), shape (Dumoulin & Hess, 2007) and contours (Pasupathy & Connor, 2001), revealing that they are represented in the intermediate stages between low- and high-level features. These studies have been instrumental in advancing our understanding of mid-level feature processing by laying down the methodological and theoretical bases of mid-level vision research. However, further progress in revealing the nature of mid-level feature processing is stalled by a set of methodological impasses, of which we highlight three.

First, since the ground truth of mid-level features is typically only available for artificial stimuli, researchers investigating ecologically more valid naturalistic stimuli often rely on model approximations or data-driven feature representations (e.g., deep neural network layer (DNN) activations). These approximations are prone to error, are noisy, and may not accurately represent human perception, especially in terms of mid-level features (see Geirhos et al., 2022).

Second, most stimulus sets focus on just one or two mid-level features, often related to shape and texture (Backus et al., 2001; Beeck et al., 2008; Freeman et al., 2013), while mid-level features form a large and diverse set, including surface- and depth-based information, requiring an integrated investigation of multiple features for a more comprehensive view. Third, the majority of stimulus sets are composed of static images despite the importance of motion processing in mid-level vision (Roe et al., 2012). This limits our understanding of how motion impacts mid-level feature processing, and leaves open whether insights from experiments using still images transfer to moving stimuli.

Here, to address these constraints we created a novel stimulus set of 3D-rendered naturalistic scenes as still images and videos with accompanying ground-truth annotations (Epic Games, 2019). This addresses the three constraints enumerated above: it provides noise- and error-less ground-truth annotations for mid-level features in naturalistic stimuli, it covers multiple mid-level features (i.e., reflectance, lighting, world normals, scene depth and skeleton position, as well as a low-level feature, edges, and a high-level feature, action identity), and it does so for both static and dynamic stimuli.

Thus equipped, we assessed the nature of mid-level feature processing in humans in two ways. First, we determined the temporal dynamics of mid-level feature representations in the human brain using electroencephalography (EEG) and linearized encoding models (Naselaris et al., 2011). Second, we determined the hierarchies of feature processing in image computable convolutional neural networks (CNNs), i.e., some of the current best models of human brain activity (Bonner & Epstein, 2018; Groen et al., 2018; Khaligh-Razavi & Kriegeskorte, 2014; Yamins & DiCarlo, 2016), to provide a computational account of mid-level feature processing (Cichy & Kaiser, 2019). Altogether, using a novel stimulus set, we provided an empirical and computational account of the temporal dynamics of mid-level features in humans during scene perception.

## 2. Materials and Methods

### 2.1 Participants

The study consisted of two experiments: an image experiment, where one set of participants viewed static images of scenes, and a video experiment, where another set of participants viewed dynamic videos of the same scenes. 15 healthy participants took part in the image experiment (mean age 23.5, *SD* = 2.58; 9 female, 6 male), and 20 healthy participants took part in the video experiment (mean age 25.15, *SD* = 4.33; 17 female, 3 male). All participants had normal or corrected-to-normal vision. All participants provided their informed consent after getting acquainted with the study protocol. The study was approved by the ethics committee of Freie Universität Berlin.

### 2.2 Stimuli and ground-truth annotations

We created the stimuli (**Figure 1A**) and ground-truth annotations (**Figure 1B**) using Unreal Engine (Epic Games, 2019). Each stimulus depicted a naturalistic scene where a character performed an action inside a room. The stimulus set consisted of scenes showing one out of 3 different characters (a woman in a white T-shirt, a man in a black T-shirt and a man in a green T-shirt), performing one of 6 different actions (arm stretching, cheering while sitting, picking up a bottle from the floor, sit-ups, standing up, and playing guitar), in one of 20 different rooms (each having a different layout and objects) and filmed from one of 4 different camera spawn points (i.e., locations where the camera is filming from approximately), resulting in a total of 1440 stimuli. The stimuli were available as 520 x 390 pixel still images and 300 ms videos of the same dimensions with a frame rate of 30 fps (i.e., containing 9 frames).

**Figure 1.**
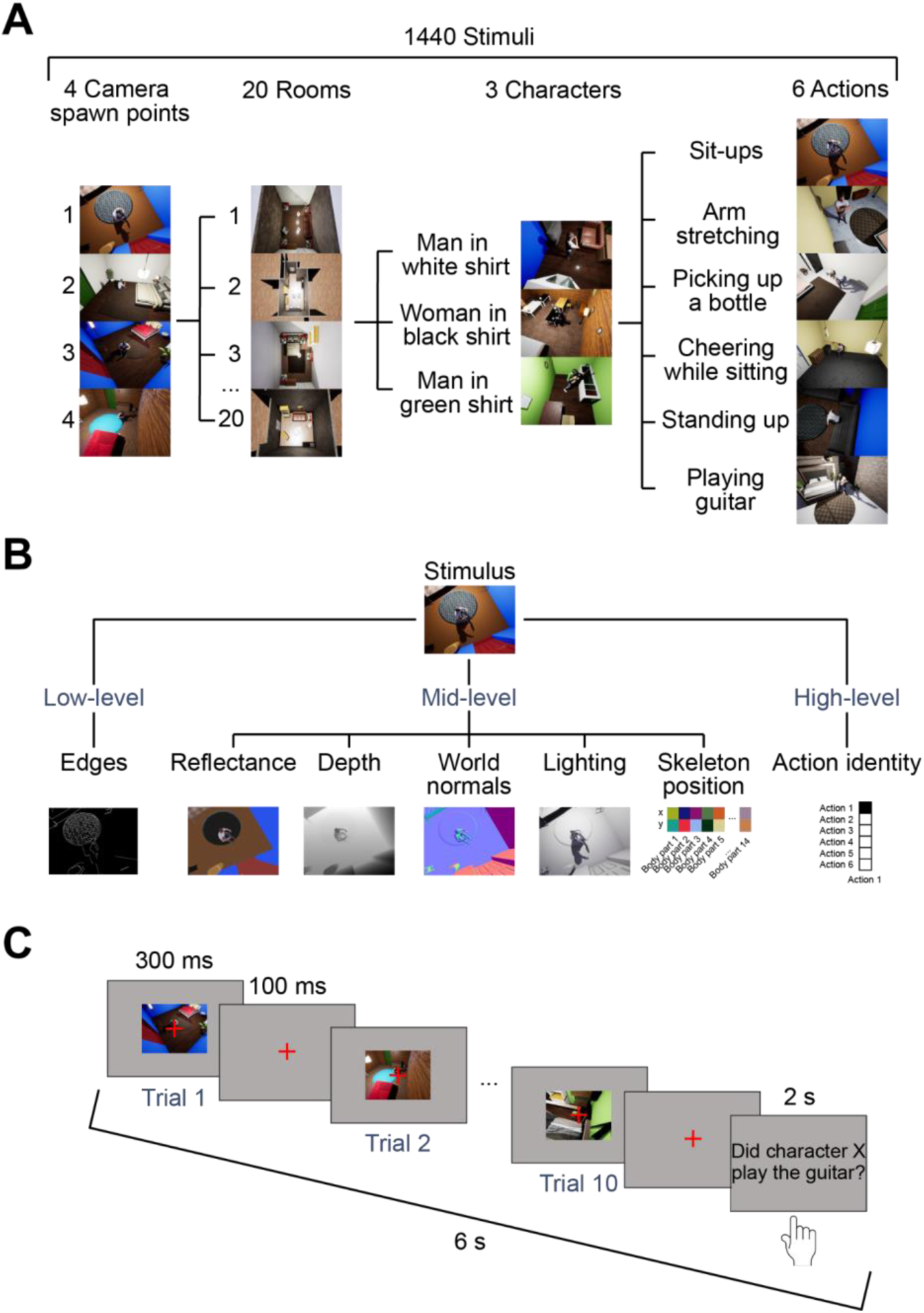
Stimuli, ground-truth annotations and paradigm. **A.** Examples of stimuli created in a game engine (Epic Games, 2019), available as images and 300 ms videos.The stimulus set contained 1440 samples, created by using 4 camera spawn points, 20 rooms, 3 characters and 6 actions. **B.** Example annotations created in the game engine for the low-, mid-, and high-level features of interest. **C.** Paradigm used in the experiment. Each trial sequence lasted 6 s, consisting of 10 trials of 400 ms each (300 ms-long stimulus presentation followed by 100 ms of interstimulus interval (ISI)) and 2 s of response time. The participants’ task was to indicate whether a specific character X (either man in white shirt, woman in black shirt or man in green shirt) was shown playing the guitar at least once in the sequence.

For every stimulus and frame (for videos), we obtained ground-truth annotations of five mid-level features - reflectance, lighting, world normals, scene depth and skeleton position - as well as for two framing features: one low-level feature, edges, and one high-level feature, action identity (**Figure 1B**). We rendered the ground-truth annotations for four of the features (reflectance, lighting, world normals and scene depth) from the game engine. The rendered annotations for the colorful features (reflectance and world normals) were of size 520 x 390 x 3, representing pixel-wise, RGB channel-wise ground-truth information for the given feature. The annotations for the grayscale features (lighting and scene depth) were of size 520 x 390.

For the remaining three features (edges, skeleton position and action identity), we obtained the annotations differently. To obtain edges, we used the Canny algorithm (Canny, 1986) as implemented in OpenCV (Bradski, 2000). This resulted in a matrix of size 520 x 390, representing pixel-wise edge information for each stimulus and frame (for videos). For skeleton position, i.e., the x and y coordinates of the character’s 14 body parts (head, neck, left upper arm, right upper arm, left lower arm, right upper arm, left hand, right hand, left thigh, right thigh, left calf, right calf, left foot, right foot), we obtained the information from the meta-data files provided by the engine, resulting in a matrix of size 14 x 2. Lastly, for action identity, i.e., the action performed by the character, we created one-hot vectors for each of the six actions and used them to encode the action information for each of the scenes. This resulted in one ground-truth annotation per stimulus and feature for images, and one ground-truth annotation per stimulus, frame and feature for videos.

Finally, for training and evaluating encoding models, the stimuli and their annotations were split into training, test and validation sets, each respectively containing 1080, 180, and 180 samples. The training set was composed of scenes spanning 15 rooms, 6 actions, 4 camera angles and 3 characters, while the test and validation sets were each composed of scenes spanning 5 rooms, 6 actions, 2 camera angles and 3 characters.

### 2.3 Paradigm and experimental design

In both image and video experiments, participants completed a rapid serial visual presentation (RSVP) task (**Figure 1C**). Each of the image and video experiments consisted of 22 self-initiated runs assigned to training, test, and validation sets (10, 10 and 2 runs, respectively). The training and test runs alternated every run, and the validation runs were added in between (one in the middle and one in the end). The training and test runs contained 54 sequences of 10 trials each, while the validation set contained 45 sequences. In total there were 5400 trials in the training set (5 trial repetitions for each of the 1080 stimuli), 5400 trials in the test set (30 trial repetitions for each of the 180 stimuli) and 900 trials in the validation set (5 trial repetitions for each of the 180 stimuli). The order of the presented stimuli was randomized for every run separately.

On each trial, participants viewed a stimulus for 300 ms (a still image in the image experiment or a 300-ms video in the video experiment) at a visual angle of 15.8° by 10.6°, which was followed by an interstimulus interval (ISI) of 100 ms showing a gray screen, totalling 4 s of presentation time for each sequence of 10 trials. We selected 300 ms as stimulus duration because it is short enough to minimize the effect of eye movements on EEG data, but long enough for the movement information to still be present in the videos. A red fixation cross was at the center of the screen throughout the stimulus presentation and the ISI to encourage participants to fixate. After these 4 s of presentation time, participants had 2 s to indicate whether they saw a target character (a woman in a black T-shirt, a man in a white T-shirt, or a man in a green T-shirt) play the guitar by pressing “J” on odd runs and “F” on even runs if they did, and the opposite key if they did not. The target character remained the same throughout each run. For train, test, and validation runs, the target characters were presented in the same repeating order: the man in the white T-shirt, the man in the green T-shirt, and the woman in the black T-shirt. Participants were also invited to blink during this period.

Prior to data collection, in order to familiarize themselves with the task, participants completed seven practice runs each containing three sequences.

The experiment was programmed using PsychToolbox (Brainard, 1997) in MATLAB (2021a).

### 2.4 EEG recording and preprocessing

During the image and video experiments, we recorded electroencephalography (EEG) data using a 64-channel actiCAP active electrode system from BrainVision with a sampling rate of 1000 Hz. The signal was amplified using the BrainAmp amplifier. The electrodes on the Easycap 64-electrode system were arranged based on the 10-10 system. The participants wore actiCAP elastic caps, connected to 64 active scalp electrodes, plus one ground (AFz) and one reference electrode (FCz). The signal was filtered online between 0.03 and 100 Hz to eliminate baseline drift and high-frequency noise.

Offline, we preprocessed the EEG data using the MNE package (1.2.2) (Gramfort et al., 2013) in Python (3.10). Preprocessing included filtering line noise with a notch filter (50 Hz), independent component analysis (ICA) on epoched video sequences to remove artifacts such as eye movements and muscle activity, low-pass filtering (25 Hz), resampling to 50 Hz (70 time points), and selecting 19 posterior (parietal and occipital) channels (i.e.,channels overlaying the visual cortex) for computational efficiency, and dividing the sequences into epochs surrounding individual stimuli. This resulted in epochs starting from 400 ms before stimulus onset to 980 ms after stimulus onset, which we baseline corrected based on a 100 ms pre-stimulus window.

To control for the different levels of noise in the electrodes, we applied multivariate noise normalization (MVNN) (Guggenmos et al., 2018). MVNN was estimated using the training set and applied to the training, test and validation sets.

At the end of preprocessing, we obtained for each trial and for every time point a pattern of 19 channel activations, which we used to perform the analyses described below.

### 2.5 Decoding

To determine and compare the signal-to-noise ratios of our image and video data, we assessed the decodability of individual scenes by performing pairwise decoding (Cichy et al., 2014; Grootswagers et al., 2017) on subject-level preprocessed EEG data. For this, we used a linear support vector machine (SVM) (Vapnik, 1995) in scikit-learn (Pedregosa et al., 2011).

We performed pairwise decoding on every pair of scenes from the test set of the encoding analysis, independently for image and video data, and for every subject and every time point. We performed the analysis on the 180 scenes from the encoding test set given that it had the largest number of trials (30).

This procedure involved two steps. First, to increase the signal-to-noise ratio for each scene, we created 6 pseudotrials from the 30 trial repetitions by averaging over randomly formed bins of 5 trials. This resulted in 6 pseudotrials per scene and therefore, 12 pseudotrials for each pairwise combination. Second, for each pairwise combination, we employed a stratified 6-fold cross-validation procedure (Cichy et al., 2014; Grootswagers et al., 2017). Using this procedure we first randomly split the 12 pseudotrials from each pairwise combination into a decoding training set and a decoding test set. We then trained the SVM on the decoding training set to classify every pair of scenes and evaluated it on the decoding test set. We obtained a symmetric matrix of pairwise decoding accuracies for all 180 x 180 pairs of scenes, of which we took the lower diagonal, flattened it and averaged across all entries, resulting in one average decoding accuracy per fold. We repeated this analysis for every fold (i.e., 6 times), using a different decoding training and test set each time, and averaged over the results across folds.

After performing this analysis on every time point, for each participant, for both image and video data, we averaged the results across participants and obtained one decoding time course for images and one for videos. We compared these two time courses by subtracting the video results from the image results, providing us with a measure of difference between the decodability, and therefore, of signal-to-noise ratio, of our two datasets.

### 2.6 Encoding

To determine when the selected low-, mid- and high-level features are processed in the brain (EEG) and whether there is a comparable processing hierarchy across the layers of convolutional neural networks (CNNs) trained on vision tasks, we trained and evaluated linearized encoding models (Naselaris et al., 2011). We first give an overview of the workflow before detailing each step. First, we preprocessed our ground-truth annotations into a suitable format for training the encoding models. Then, for CNNs, we extracted and preprocessed activations at different layers throughout the networks. On this basis, separately for EEG and CNNs and for images and videos, we used ridge regression (Hastie et al., 2009) to train encoding models to predict the EEG signal at every time point, or CNN activations at every layer, from our low-, mid- and high-level feature ground-truth annotations. We then assessed the accuracy of the encoding models by correlating between the predicted and true data, which revealed when low-, mid-, and high-level features are represented in humans and CNNs. Lastly, we collected the encoding peak latencies for EEG and peak layers for CNNs for all features and correlated them, revealing similarities and dissimilarities in the processing hierarchy of low-, mid- and high-level features in humans and CNNs.

#### 2.6.1 Preparing the ground-truth annotations

To put the image ground-truth annotations of the low-, mid- and high-level features into a format suitable for the encoding analysis, we prepared them by applying two preprocessing steps (**Figure 2A**). First, for each feature, except skeleton position and action identity, and each stimulus in the training, validation and test sets, we flattened the annotations along the width, height, and for colorful features, RGB dimensions. For skeleton position we flattened the annotations across x/y coordinates to transform them into 1D vectors, and action identity was already present in a 1D format. Second, to improve the computational efficiency of the encoding analysis, we reduced the dimensionality of the annotations for the higher-dimensional features, i.e., edges, reflectance, lighting, world normals and scene depth. For this, we employed a principal component analysis (PCA) with a linear kernel and 100 components using scikit-learn in Python (3.10). PCA was not employed on the two lower-dimensionality features, i.e., skeleton position and action identity, as they already contained less than 100 components (28 for skeleton position and 6 for action identity).

**Figure 2.**
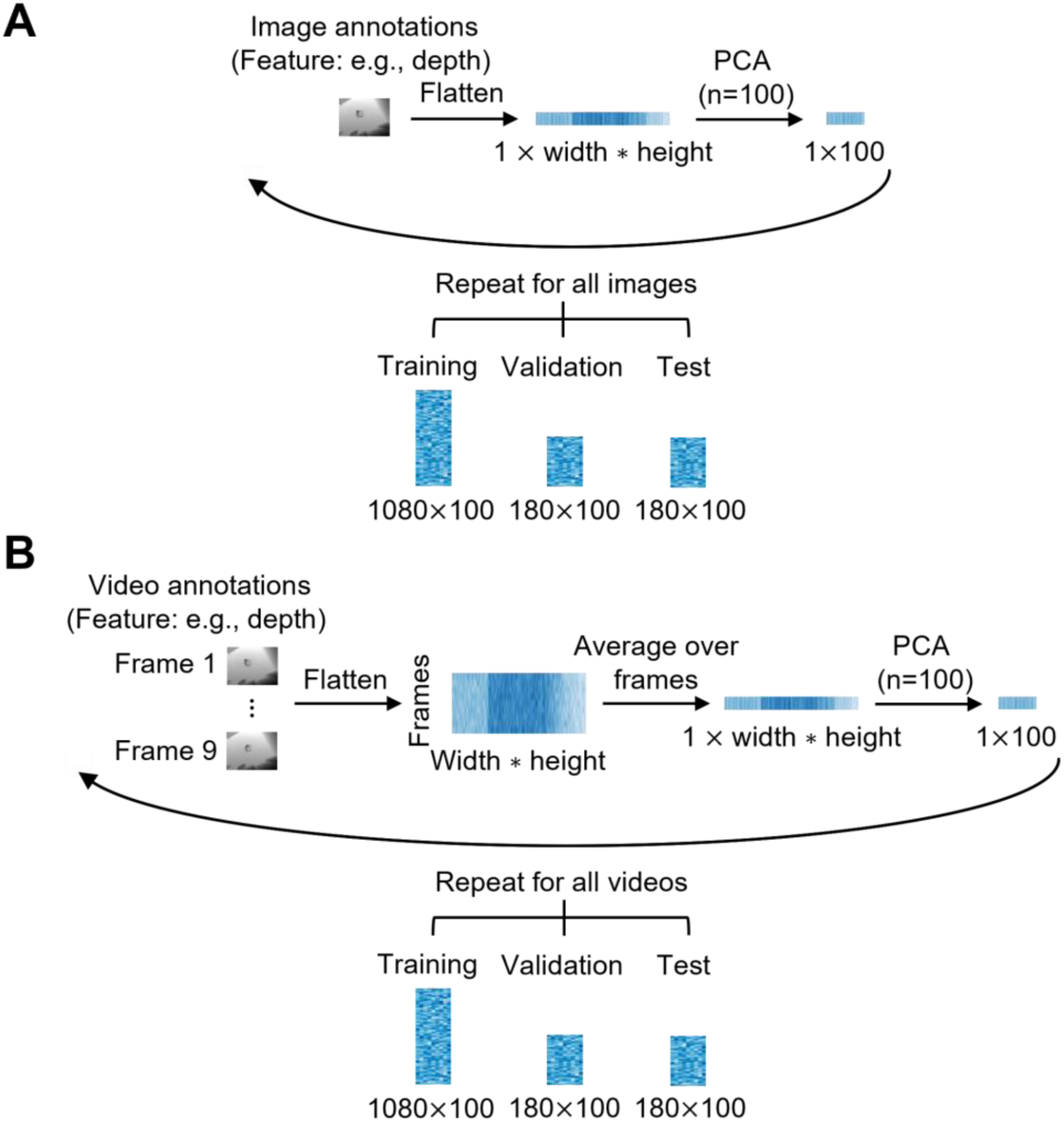
Preparation of ground-truth annotations. **A.** Image annotations. For each feature separately, the ground-truth annotation for every image was 1) flattened across the width, height and if applicable, RGB dimensions, and 2) reduced in dimensionality via PCA (100 components) for the high-dimensional features. Repeating the procedure over all images resulted in vectors of size 100 for every image from the training set (1080 samples), validation set (180 samples) and test set (180 samples). **B.** Video annotations. For each feature separately, the ground-truth annotations for all 9 frames were 1) flattened across the width, height and if applicable, RGB dimensions, 2) averaged over the 9 frames, and 3) reduced in dimensionality via PCA (100 components) for the higher-dimensionality features. Repeating the procedure for all videos resulted in vectors of size 100 for every video from the training set (1080 samples), validation set (180 samples) and test set (180 samples).

To prepare the video ground-truth annotations (**Figure 2B**), we applied three preprocessing steps. First, we flattened the annotations across the width, height and when applicable, RGB dimensions for each feature (except for skeleton position and action identity), stimulus and video frame, and across x/y coordinates for skeleton position. Second, for each feature and stimulus, we averaged over the frames. Third, for each stimulus, we applied PCA with 100 components on the averaged annotations for the five higher-dimensional features.

The image and video annotations for these low-, mid-, and high-level features were then used to train encoding models and predict EEG and CNN data.

#### 2.6.2 Extracting and preparing the CNN activations

To train and evaluate encoding models on ground-truth annotations and CNNs, we first extracted CNN activations and then in a second step prepared them for the encoding analysis.

The details of the first step were as follows. We first extracted hidden layer activations from two CNNs: one trained and evaluated on scene classification using images as inputs, referred to as image CNN, and one trained and evaluated on action classification using videos as inputs, referred to as video CNN.

For the image CNN, we used an instance of ResNet-18 (He et al., 2015) pre-trained on Places365-Standard (Zhou et al., 2018) (https://github.com/CSAILVision/places365), a dataset of more than 1.8 million scene images across 365 scene categories. We extracted activations from its hidden layers in response to all the images from our stimulus set. Before extracting the activations, we preprocessed the images to put them in a suitable format for the network. For this, we center-cropped the image frames from the encoding training, test and validation sets, resized them to 224 x 224 pixels and normalized them. Afterwards, we fed the preprocessed images to the network and extracted the activations from the last layers of eight residual blocks (referred to as layer 1.0, layer 1.1, layer 2.0, layer 2.1, layer 3.0, layer 3.1, layer 4.0 and layer 4.1) using Torch FX in PyTorch in Python (3.10), yielding activations from eight layers throughout the network hierarchy for the training, validation and test sets of images.

For the video CNN, we used an instance of 3D ResNet-18, a network that employs 3D convolutions on the spatiotemporal video volume (Tran et al., 2018), from the TorchVision video class. It was pre-trained on the Kinetics400 dataset comprising up to 650,000 videos that cover 400 human actions (Kay et al., 2017). We extracted activations from the same eight hidden layers as from the image CNN, in response to all the videos from our stimulus set. Before extracting the activations, we preprocessed the video frames: we rescaled them to 128 x 171 pixels, then center-cropped and resized them to 112 x 112 pixels, and finally normalized them. Then, we fed these preprocessed frames to the CNN, and extracted the activations from the last layers of its eight residual blocks (the same ones as in the image CNN) using Torch FX in PyTorch in Python (3.10), obtaining activations from eight layers in response to the videos from our training, validation and test sets of videos.

The second step was to prepare the extracted CNN activations for encoding. We flattened the activations for each layer of the image and video CNN across all feature map dimensions, obtaining a 2D matrix of size number of images x number of CNN features. To improve computational efficiency, we reduced the dimensionality of the flattened activations by applying PCA with the number of components that explained at least 90% of the variance (see **Supplementary Table 1** for the numbers of components), separately for each layer of each image and video CNN separately. This yielded CNN activations for eight layers for all stimuli from the training, validation and test sets, suitably formatted for further encoding analyses.

#### 2.6.3. Predicting the EEG signal from ground-truth annotations

To determine when low-, mid- and high-level features are processed in the brain, we trained and evaluated linearizing encoding models (Naselaris et al., 2011) (**Figure 3A**). We conducted all procedures separately for images and videos, for each subject, feature, EEG channel and time point.

**Figure 3.**
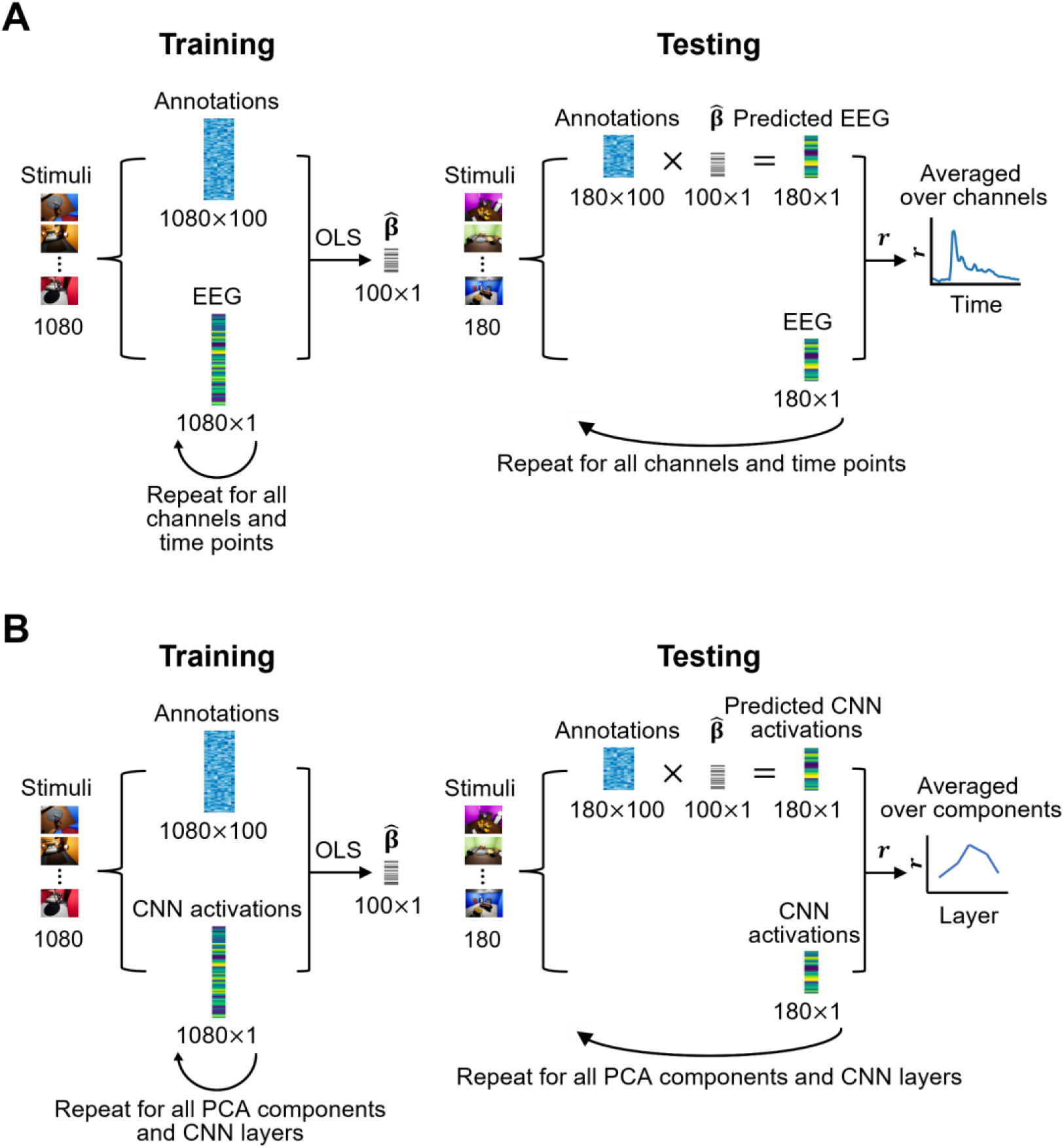
Encoding analysis pipeline. **A.** EEG encoding pipeline for one feature. The training set of stimuli and annotations is used to estimate the regression weights **β** of the regression model, where the annotations predict EEG data at every channel and time point. Afterwards, the test set of stimuli and annotations is used to predict EEG data from annotations with the estimated regression weights at every channel and time point. The predicted EEG data is then correlated with true EEG data to assess the accuracy of the encoding model at every time point and channel, and the final time course depicts the average correlation over channels. **B.** CNN encoding pipeline for one feature. The training set of stimuli and annotations is used to estimate the regression weights **β** of the regression model, where the annotations predict CNN activations at every PCA component and layer. Afterwards, the test set of stimuli and annotations is used to predict CNN activations from annotations with the estimated regression weights at every PCA component and layer. The predicted CNN activations are then correlated with true CNN activations to assess the accuracy of the encoding model at every time layer and PCA component, and the final layer-wise plot depicts the average correlation over PCA components.

In short, we first trained models using multivariate linear ridge regression (L2 regularization) with an optimized *λ* value, and then used these models to predict EEG data from the preprocessed ground-truth annotations, separately.

In detail, the first part of the procedure was hyperparameter *λ* optimization and involved three steps. First, we estimated the *β* regression weights by using the ordinary least squares (OLS) method as described in (Hastie, 2020). The hyperparameter space spanned values in the logarithmic range from 10^−5^ to 10^10^. Second, we assessed the accuracy of the regression model in predicting the EEG signal on the validation test using Pearson’s correlation. Third, we determined the optimal *λ* based on the encoding accuracy averaged across channels and time points. The optimal value of *λ* for each feature and subject was then used to perform the rest of the encoding analysis.

To conduct the second part of the procedure, using the optimized lambda from the first part, we trained a regression model on the training set to predict the EEG signals using the annotations as predictors. We used the model to predict the EEG signal of the test set from the annotations. For every subject and every feature, we correlated the predicted EEG data for each channel with the true EEG data for each channel from the test set. We averaged over channels and subjects, performed the analysis on image and video data separately, and obtained one time course for every low-, mid-, and high-level feature for images and one for videos.

#### 2.6.4. Predicting CNN activations from ground-truth annotations

To determine the relationship between CNN layers and stimulus low-, mid- and high-level features, we trained and evaluated linearizing encoding models to predict the dimensionality-reduced activations, i.e., the principal components, from our ground-truth annotations (**Figure 3B**). For this, we trained ridge-regression models (akin to the EEG encoding analysis) on flattened and dimensionality-reduced image and video CNN activations separately, for every feature, layer, and principal component, and evaluated them by predicting the respective test set activations and correlating the predicted and true activations for every feature, layer and principal component. In order to account for the different levels of variance explained by every component, we then calculated, for every feature and layer, the weighted average of the component-specific correlations. In detail, we weighed each component’s correlation by its amount of explained variance, summed the weighted correlations and divided the result by the total variance explained. We implemented this approach to assign greater weights to components that contribute more to the overall variance, ensuring that the final correlation results reflect the most informative components rather than being equally influenced by all components. This resulted in layer-wise correlations characterizing the processing of low-, mid-, and high-level features in image and video CNNs.

#### 2.6.5 EEG noise ceiling

To estimate the theoretical maximum encoding correlation results given the level of noise in our EEG data, we calculated the lower and upper bound of the noise ceiling (Gifford et al., 2022). We computed noise ceiling estimates for every subject and for image and video data separately. To calculate the lower bound of the noise ceiling, we split all the trials into two equal groups and correlated the preprocessed channel-wise, time point-wise EEG data between the two groups. To calculate the upper bound of the noise ceiling, we correlated between a group containing half of all trials and a group containing all trials. We repeated the calculation of the lower and upper noise ceilings 100 times to permute the grouping of the trials each time and averaged over the permutations and subjects to obtain the final lower and upper noise ceilings.

#### 2.6.6 Correlation between EEG peak latencies and CNN peak layers

To determine whether low-, mid- and high-level features are processed in a similar hierarchy between humans and CNNs, we correlated between the EEG peak latencies and CCN peak layers. For this, we first collected the encoding peak latencies for every one of the seven features for the EEG and the peak layers from the CNN analyses separately, for images and videos separately. This resulted in four 7×1 vectors. Then, for images and videos separately, we calculated the Spearman’s correlation between the EEG and CNN vectors, revealing the relationship between the processing hierarchies of humans and CNNs.

### 2.7 Statistical analysis

To determine the statistical significance of the decoding and encoding results, we performed non-parametric statistical tests. This involved three steps, performed separately for each EEG time point and CNN layer. First, we used the sign-test to create 10 000 permutation samples. For EEG decoding and encoding, this involved multiplying 10 000 times the subject-level decoding or encoding results by a random vector of 1s and −1s. For CNN encoding, this meant multiplying 10 000 times the results for each principal component by a random vector of 1s and −1s. Second, we obtained the p-value of the true data by calculating the rank of its mean with respect to the distribution of the permutation samples. Third, we controlled for multiple comparisons by adjusting the p-values of the original data for their inflated false discovery rate (FDR) using the Benjamini-Hochberg procedure (Benjamini & Hochberg, 1995) with ⍺=0.05 (two-tailed).

Additionally, we calculated the bootstrapped (with replacement) 95% confidence intervals (CIs) of the EEG decoding and EEG encoding accuracies to obtain an approximation of the sampling distribution. The 95% CIs computed from this distribution are expected to contain the true population mean at 95% of the time., To approximate the distribution, separately for each of the EEG decoding and EEG feature-specific image and video encoding time courses, we bootstrapped the subject-level accuracies 10 000 times and calculated the 95% confidence intervals (short CIs, i.e., the values at the 2.5th and the 97.5 percentiles) of the bootstrapped distribution. To have an analogous measure of the range of possible results in CNNs, we also calculated the CI intervals for the CNN encoding time courses. For this, we sampled with replacement the results for the principal components of each layer 10 000 times and calculated the 95% CIs of the bootstrapped distribution of the encoding accuracies.

Moreover, we calculated the bootstrapped 95% CIs for the peak latencies of EEG and peak layers of the CNN. We used these CIs to 1) estimate the uncertainty around our sample peak latencies and layers and to 2) test whether the differences between the encoding peak latencies and layers for the different features were statistically significant. To address the latter, for each pairwise combination of features from the image and video results separately, we created the bootstrapped distribution for the difference between peaks, and whenever the 95% CI for the difference between two conditions contained 0, the difference was deemed non-significant.

To determine the significance of the EEG peak latency and CNN peak layer correlation, we used a three-step approach. First, we performed the sign-test to create 10 000 permutation samples of the CNN encoding results. Second, we calculated the mean of each sample across principal components, for each layer and feature separately. Third, for each sample, we calculated the peak layers for all features and correlated them with the EEG peak latencies. This created a distribution of sample correlations based on which we determined whether the original correlation was significant or not (i.e., if its p-value was smaller than 0.05).

## 3. Results

### 3.1 Decoding images and videos from EEG yields comparable results

To determine whether the quality of the neural signal is comparable for image and video data, as video data might be noisier due to a stronger presence of eye movements (Böhme et al., 2006), we performed time-resolved pairwise decoding (Cichy et al., 2014; Grootswagers et al., 2017) on the EEG data collected during image and video presentation and compared the decoding time courses.

We observed similar results for image and video decoding (**Figure 4**). For both, we observed an onset of decoding at 60 ms after stimulus onset (p<0.05, FDR-corrected), meaning that individual scenes were distinguishable in the brain starting from then, for both datasets. In both cases, decoding was significant (p<0.05) throughout most of the trial and peaked at 100 ms ([95% CI]: images: [100 ms, 100 ms], videos: [100 ms, 120 ms]) after stimulus onset, showing that individual scenes were best distinguishable at that time, for both images and videos. The difference wave between image and video decoding was not significant at any time point, further suggesting a similar quality of the signal between image and video EEG data. Overall, the decoding results show that scene information is as well decodable from image as from video EEG data, suggesting that the quality of the signal is comparable between the two datasets. This indicates that the two datasets can be further analyzed using the equivalent methods, and that differences observed in the results are not trivially due to data quality differences.

**Figure 4.**
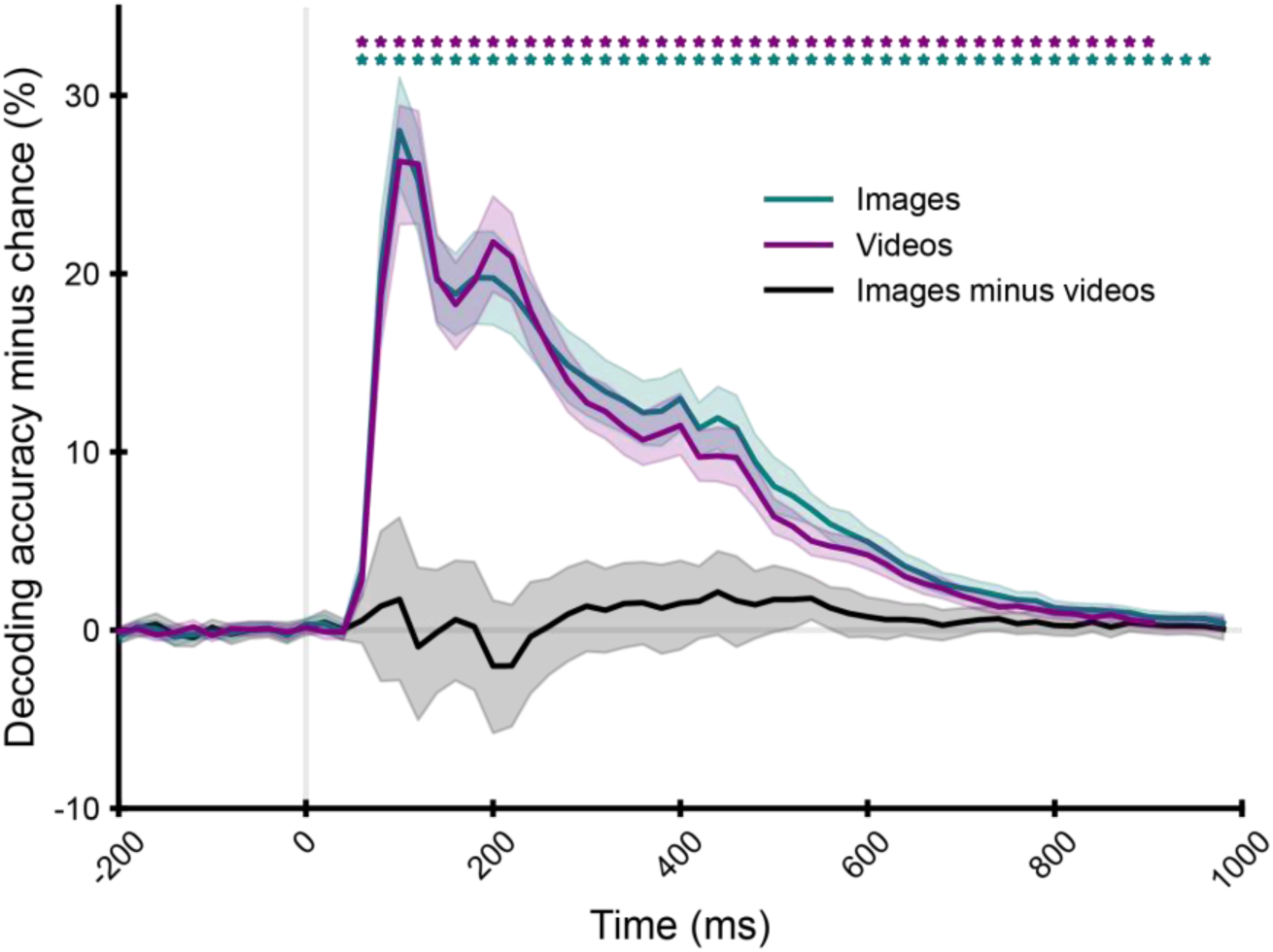
Decoding scene identity from images and videos. Averaged pairwise decoding results for individual scene images (15 subjects, green curve) and videos (20 subjects, purple curve). The difference curve (images minus videos) is shown in black. We indicated the significant time points (two-tailed, p<0.05, FDR-corrected) with stars above the time course, the 95% confidence intervals with shaded areas around the curves, the stimulus onset with a vertical gray line and the decoding chance level with a horizontal gray line.

### 3.2 Mid-level features are most strongly encoded in the brain between ∼100 ms and ∼250 ms post-stimulus

To determine the time course of mid-level feature processing in the brain with respect to low- and high-level features, we trained and evaluated encoding models (Naselaris et al., 2011) that predicted EEG data from ground-truth annotations of one low-level feature (edges), five mid-level features (reflectance, lighting, world normals, scene depth and skeleton position), and one high-level feature (action). We hypothesized that the mid-level features would be processed between the low- and high-level features, as would be expected following the classical hierarchical view of feed-forward scene processing (Groen et al., 2013; J. M. Henderson & Hollingworth, 1999). We performed the encoding analysis on image and video data separately, and compared the results to determine potential differences in feature processing.

We made three key observations. First, for both image and video data, the ground-truth annotations for all low-, mid- and high-level features predicted the EEG data significantly from ∼60 ms on (p<0.05, FDR-corrected) throughout most of the trial (**Figure 5A** and **B**, left). This demonstrates that the annotations across all feature complexities relate to the neural scene representations.

**Figure 5.**
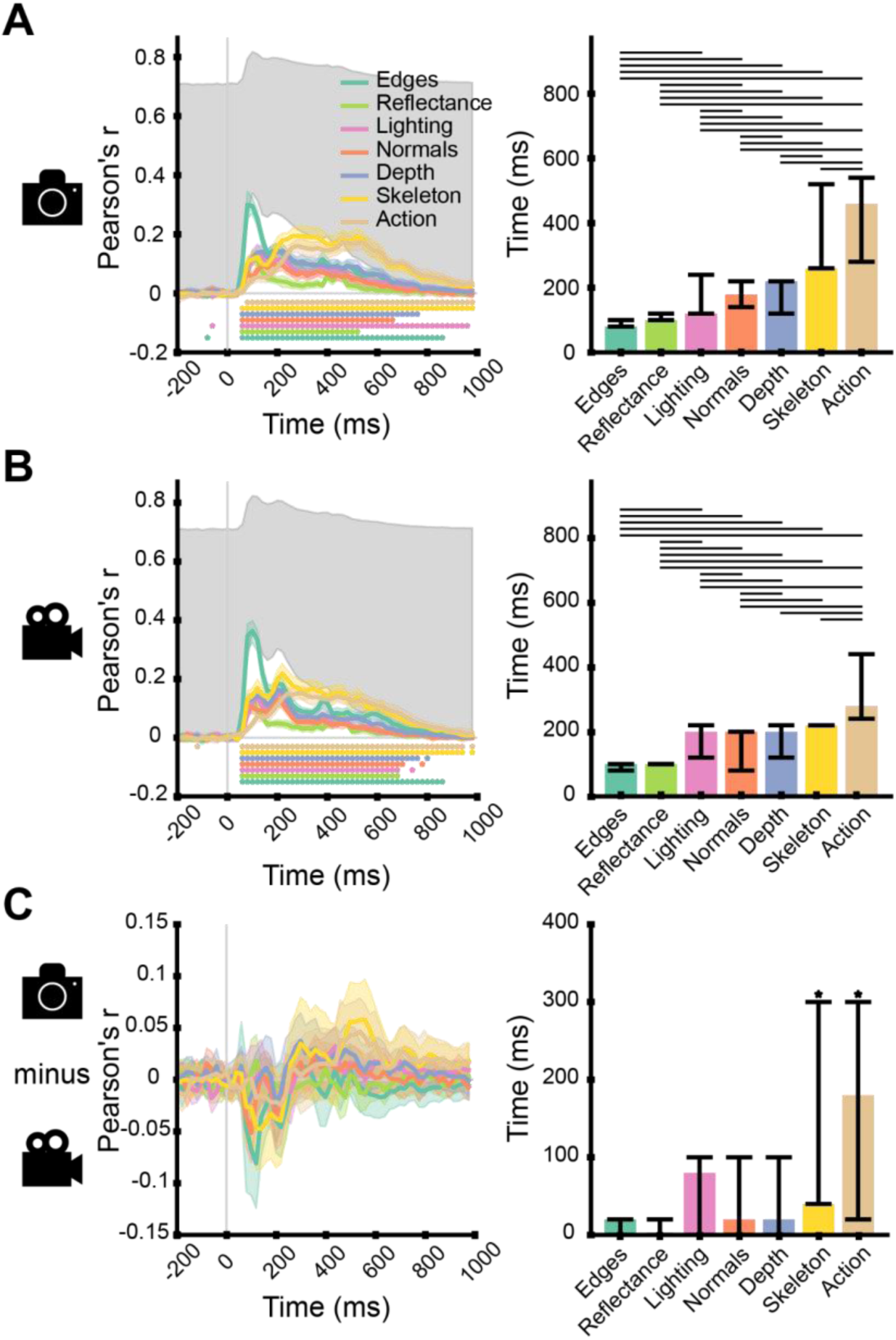
Predicting low-, mid-, and high-level feature EEG representations in scene images and videos from their ground-truth annotations. Encoding results for EEG data collected during the viewing of (**A**) images (15 subjects), (**B**) videos (20 subjects) and **(C)** their difference. Left panel: the time course of one low-, five mid-, and one high-level feature representations. We indicated the significant time points with stars (two-tailed, p<0.05, FDR-corrected), the 95% confidence intervals with shaded areas around the curves, the stimulus onset with a vertical gray line, the chance level with a horizontal gray line and the noise ceiling with a shaded gray area. Right panel: peak latencies of the features. In **C**), the right plot represents the differences in peak latencies between images and videos. We indicated the 95% confidence intervals with vertical error bars, the significance (p<0.05) between feature peak latencies with horizontal bars, and the significant peak latency differences with stars above the error bars.

Second, we observed that the encoding accuracy for mid-level features peaked between ∼100 ms and ∼250 ms after stimulus onset, for both images and videos (**Figure 5A** and **B**, right). Of the mid-level features, for both images and videos, the encoding accuracy peaked first for reflectance and last for skeleton position, while the results for lighting, world normals and scene depth were in-between them and peaked close to one another (see **SupplementaryTable 2** for statistical details). For both images and videos, the results for all mid-level features except for reflectance peaked significantly after the low-level feature, i.e., edges (on average, 90 ms later, significant differences indicated by lines above bar plots), and significantly before the high-level feature, i.e., action identity (on average, 100 ms earlier). This suggests that mid-level features are processed between our selected low- and high-level features, as predicted by the hierarchical view of scene perception (Groen et al., 2017; J. M. Henderson & Hollingworth, 1999).

Third, there were no significant differences in the encoding accuracy between images and videos for any feature (**Figure 5C**, left). However, the encoding peak latency was significantly different between images and videos for one mid-level feature, skeleton position, and the high-level feature, action identity (**Figure 5C**, right). Indeed, skeleton position and action identity peaked significantly later in images than in videos (skeleton position difference: 40 ms, 95% CI: [40, 300]; action identity difference: 180 ms, 95% CI: [20, 300]). This suggests that the dynamic changes in videos accelerate resolving information related to biological motion (Johansson, 1973), such as action identity and skeleton position.

Overall, our results indicate that mid-level features are processed between ∼100 ms and ∼250 ms after stimulus onset, i.e., between low- and high-level features, for both images and videos, and that features related to biological motion such as skeleton position and action identity are processed more quickly during video presentation than during image presentation.

### 3.3 Mid-level features are processed in CNNs throughout the layer hierarchy

To determine the layer-wise representations of mid-level features in CNNs and whether CNNs represent mid-level features similarly to humans with respect to low- and high-level features, we trained and evaluated encoding models that predicted CNN layer activations during stimulus presentation from ground-truth annotations, separately for images and videos. For this, we selected one image CNN pre-trained on scene classification with images (ResNet-18) (He et al., 2015; Zhou et al., 2018) and one video CNN pre-trained on action classification with videos (3D ResNet-18) (He et al., 2015; Tran et al., 2018) and collected their activations from eight layers in response to the image and video stimuli, respectively. We then trained encoding models to predict their activations to the training set stimuli from ground-truth annotations. Then, for each network, layer, and feature, we applied the trained encoding models to predict the activations to the test set stimuli from ground-truth annotations. Lastly, we calculated the encoding accuracy by correlating the predicted and the true activations for every network, layer and feature, thereby characterizing the layer-wise processing of low-, mid- and high-level features for image and video CNNs.

We made three main observations.

First, for both image and video CNNs, the ground-truth annotations predicted significantly (p<0.05, FDR-corrected) the activations for all features and layers (**Figure 6A** and **B**, left, significance indicated by stars). This shows that the (employed) ground-truth annotations are suitable for modeling the processing of low-, mid- and high-level features in CNNs, thus warranting further investigation.

**Figure 6.**
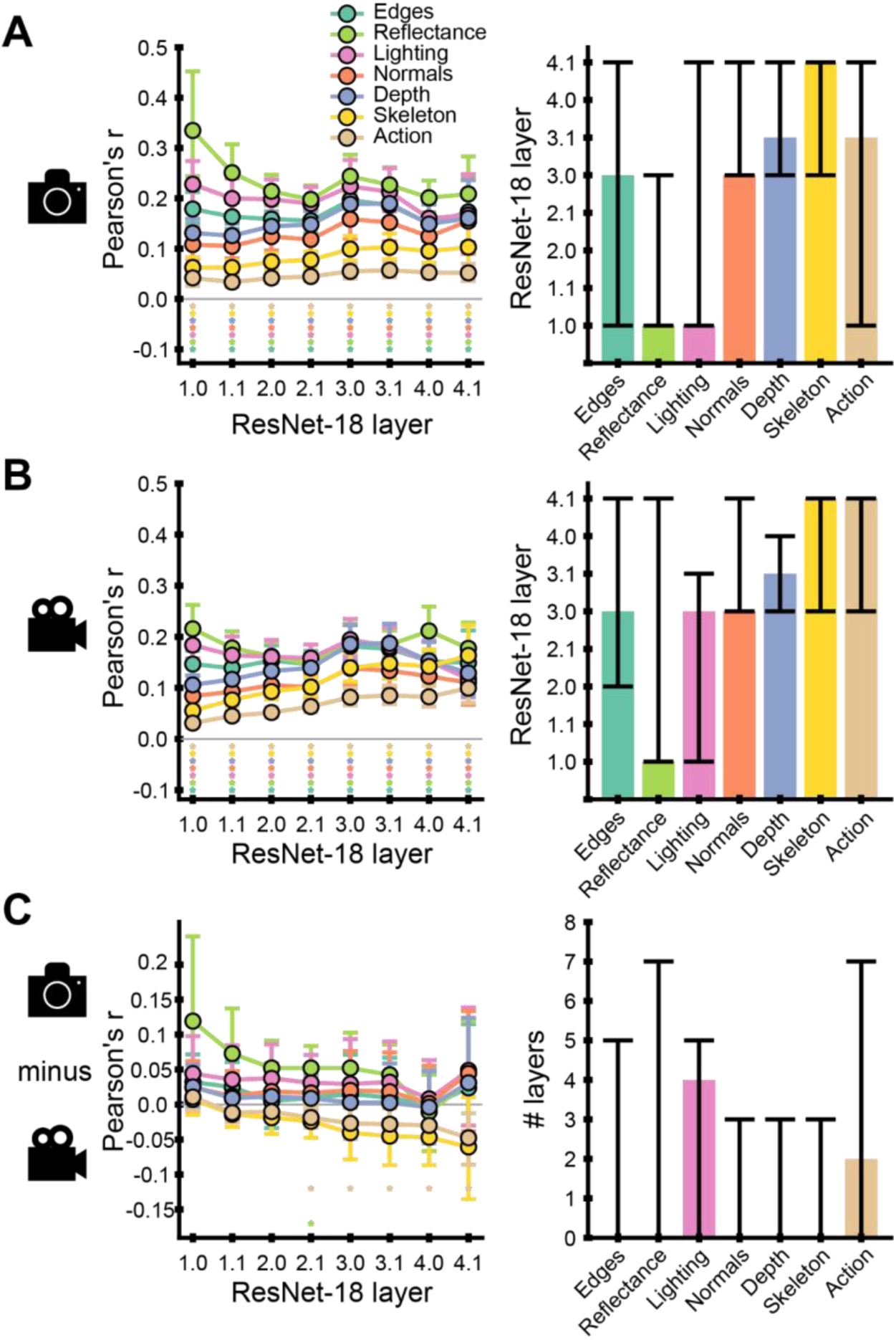
Predicting low-, mid-, and high-level feature CNN representations in scene images and videos from their ground-truth annotations. Encoding results for CNN activations collected during the presentation of (**A**) images to ResNet-18 trained on scene classification, (**B**) videos to 3D ResNet-18 trained on action classification, and **(C)** their difference. Left panel: encoding accuracy for one low-, five mid-, and one high-level feature representations, for eight CNN layers (named 1.0, 1.1, 2.0, 2.1, 3.0, 3.1, 4.0 and 4.1). We indicated the peak layers with dashed lines, the significant layers with stars (two-tailed, p<0.05, FDR-corrected) and the 95% confidence intervals with shaded areas around the curves. Right panel: peak layers of the encoding accuracies for the seven features. In **C**), the right plot represents the differences in peak layers between images and videos. We indicated the 95% confidence intervals with vertical error bars and the significance (p<0.05) between feature peak layers with horizontal bars.

Second, mid-level features were best predicted by ground-truth annotations at various layers from the first to the penultimate one, on average at the fourth layer (layer 2.1) in images and fifth layer (layer 3.0) in videos (**Figure 6A** and **B**, right, see **Supplementary Table 2** for peak latencies and CIs). This suggests that mid-level features are processed throughout the hierarchy of CNNs, and most strongly on average in intermediate processing stages.

Third, we observed significant (p<0.05, FDR-corrected) differences between the image and video CNN encoding accuracies for action identity from the fourth to the eighth layers (**Figure 6C**, left). For action identity, the ground-truth annotations better predicted the activations of the video CNN than of the image CNN for all eight layers, demonstrating that CNNs trained on action classification with videos represent action-related features more strongly than CNNs trained on scene classification with images.

To summarize, our encoding analysis on activations from CNNs trained on scene and action classification revealed that mid-level features are processed most strongly at intermediate CNN, with significant differences in encoding accuracy for action identity.

### 3.4 The processing hierarchy of mid-level features correlates between humans and CNNs

To determine whether humans and CNNs process low-, mid- and high-level features via a similar hierarchy, assuming an analogy between processing time in the brain and layer depth in CNNs (Cichy et al., 2016), we compared qualitatively and quantitatively (using Spearman’s correlation) between the encoding peak latencies in EEG and peak layers in the CNNs for all features, for images and videos separately.

We first observed qualitatively that ground-truth annotations predicted the CNN activations for mid-level features, but not low- or high-level features, in a similar order as in the EEG results, for both images and videos. Indeed, in both humans and CNNs, the earliest represented mid-level features were reflectance and lighting, followed by world normals, then scene depth, and finally, skeleton position. However, the low-level feature (edges) was predicted in humans before any mid-level feature, while in CNNs, it was predicted between mid-level features. The high-level feature (action) was represented in humans after all mid-level features, while in CNNs, it was represented at the same time as some mid-level features. Therefore, unlike in EEG, where edges were processed first and action identity was processed last, in CNNs, edges and action were processed synchronously with mid-level features.

We ascertained these observations quantitatively, for both images (**Figure 7A**) and videos (**Figure 7B**), the correlation between EEG and CNN peak latencies was significant (images: **ρ*=*0.77, p=0.0318; videos: **ρ**=0.88, p=0.0094). However, we note that this correlation is driven by the mid-level features: low- and high-level features do not follow the same processing order in humans and CNNs, which is further confirmed by the correlation results for the analysis using only mid-level features (images: **ρ*=*0.98, p=0.0044, videos: **ρ*=*0.92, p=0.0160).

**Figure 7.**
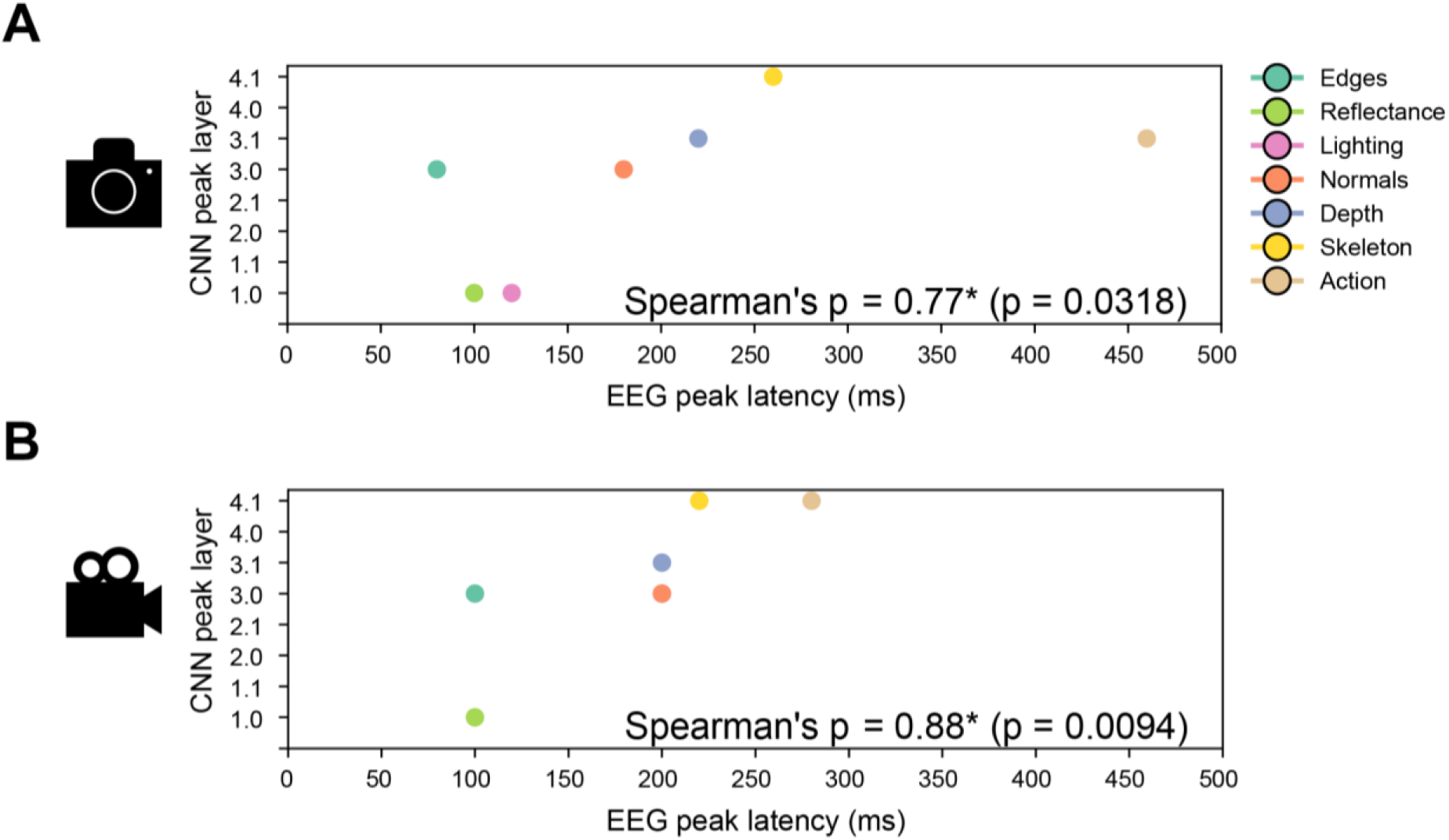
Correlating EEG encoding peak latencies and CNN encoding peak layers. Results for the Spearman’s correlation between the EEG and CNN encoding peak latencies for the analysis on (**A**) images and (**B**) videos. We indicated the Spearman’s coefficient value and its significance (p<0.05).

Overall, these results suggest that mid-level features, but not low- or high-level features are processed with a similar hierarchy in humans and artificial neural networks, both during the viewing of static images and dynamic videos.

## 4. Discussion

### Summary of results

In this study, we introduced a novel stimulus set of naturalistic images and videos with ground-truth annotations to assess the neural processing of surface-based mid-level features in humans, both empirically and via modelling. We report two main findings. First, assessing the time course of visual processing, we found that mid-level features are best represented in the brain between ∼100 and ∼250 ms post-stimulus, between low-(∼90 ms) and high-level (∼370 ms) features, with similar timing in images and videos, except for the motion-related feature skeleton position, which appeared later in images. Second, we observed that scene and action CNNs process mid-level, but not low- or high-level features, with a similar hierarchical order of processing as humans.

### Mid-level features are represented midway between low- and high-level features in humans

Leveraging a novel stimulus set with ground-truth annotations, we showed that mid-level features were best represented between ∼100 ms and ∼250 ms post-stimulus. This is consistent with previous reports that mid-level feature information can be extracted from objects between 100 and 400 ms (Grootswagers et al., 2019; Wang et al., 2022) and that outline shape and object texture are represented in the brain between 150 and 250 ms (Proklova et al., 2019). Given that low-level information such as edges and luminance is processed in the cortex as early as at 80 ms (Di Russo et al., 2002; Groen et al., 2017), and high-level semantic information such as animacy (Carlson et al., 2013; Cichy et al., 2014), spatial coherence (Groen et al., 2013) and action identity (Ge et al., 2019) is best represented at 200-400 ms and later, this places our mid-level feature representations right between low- and high-level features, consistent with a role in bridging low- and high-level vision.

We take our findings to be facilitated by three advances of our experimental approach. First, using a stimulus set with ground-truth annotations allowed us to link stimulus features and time courses of neural processing particularly strongly. Most previous studies relied on model approximations of the ground-truth such as Taskonomy outputs (Zamir et al., 2018) or on data-driven CNN activations (Kubilius et al., 2016) that contain noise and error and therefore warrant only a weaker link, capturing human processing less accurately (Geirhos et al., 2022). Moreover, CNN activations across layers approximately capture feature complexity (Cichy et al., 2016; Eickenberg et al., 2017; Güçlü & van Gerven, 2017), but are not feature specific, and thus, intermediate layers only indirectly capture human mid-level representations. In contrast, ground-truth annotations address these two shortcomings by depicting the noiseless true state and by directly assessing specific mid-level features.

Second, our stimulus set was annotated for multiple mid-level features, allowing us to capture the diverse nature of mid-level features more comprehensively than studies assessing fewer features (Beeck et al., 2008, 2008; Jagadeesh & Gardner, 2022; Yue et al., 2020), and on level ground in a unified experimental setting. In particular, we focused on surface-based features, which lack research compared to better understood features such as shape or texture (M. M. Henderson et al., 2023; Long et al., 2018; Papale et al., 2020). Our results thus add to the theory of human visual processing by showing that surface-based mid-level features are also processed in the intermediate stages.

Third, our approach characterized the mid-level feature processing in both still images and dynamic videos, allowing us to identify similarities and differences across those processing formats. We showed that in videos, skeleton position was best represented earlier than for images, which points to a quicker and more efficient resolution of action-related information for dynamic stimuli. We speculate that the motion in videos speeds up the processing of biological motion: for instance, it is easier to determine the identity of an action when viewing a video of the action than when viewing a still image of the action (Girish et al., 2020; Johansson, 1973). For the remaining mid-level features, the timing was equivalent for images and videos, showing the robustness and generalizability of our results across stimulus modalities.

### Mid-level features, but not low- or high-level features, are processed via a similar hierarchy in humans and CNNs

We compared the hierarchies of feature processing of humans with those of image-computable models to determine their suitability as brain models in this regard. As a proof of concept, we determined the predictivity of one scene- and one action-trained CNN of image and video EEG results, respectively. We made two observations.

First, we observed that mid-level feature processing follows a similar hierarchy in CNNs and humans, reinforcing prior evidence of hierarchical correspondences (Cichy et al., 2016; Guclu & van Gerven, 2015; Khaligh-Razavi & Kriegeskorte, 2014; Lahner et al., 2024). This supports previously observed similarities in the nature and processing stage of mid-level representations in CNNs and humans: for instance, (Kubilius et al., 2016) reported correlations between intermediate object-trained CNN layer activations and human perceptual similarity judgments of shape, which in turn correlated with neural activity in V3 and lateral occipital cortex LOC (Beeck et al., 2008). Similarly, (Wallis et al., 2017) showed that humans accurately classified textures generated by CNN, represented in intermediate CNN layers and macaque area V2 (Laskar et al., 2018).

Second, we found that there is a deviation in hierarchical correspondence in terms of our framing features. For one,we observed that edge processing occurs at different latencies in humans and CNNs, which we attribute to our choice of edge algorithm, canny edges. While early CNN layers are often described as edge detectors (LeCun et al., 2015; Zhou et al., 2018), they do not implement the Canny algorithm to extract edge information. On the other hand, canny edge outputs resemble silhouettes and likely engage shape-extracting mechanisms later in the CNN hierarchy. Since they share similarities with human drawings (Borji & Itti, 2014), which are processed early in humans (Singer et al., 2023) but in intermediate CNN layers (Singer et al., 2022), this latency difference between CNNs and humans is plausible. Moreover, we observed that action is represented at different stages of processing in humans and CNNs. Unlike in humans, where action is processed at later stages, we speculate that in CNNs, action is represented in intermediate-to-late layers (Peng et al., 2024), possibly due to its reliance on shape information rather than on top-down, experience-driven features. These observations align with previous studies reporting a lack of hierarchical correspondence between humans and CNNs (Sexton & Love, 2022; Xu & Vaziri-Pashkam, 2020), emphasizing the need for further research to ascertain whether humans and machines process visual input in similar stages.

### Conclusion

Using a novel approach leveraging a naturalistic stimulus set with ground-truth annotations, we revealed that mid-level features are processed after low- and before high-level features in the human brain, suggesting a bridging role. Comparing human and image-computable CNN processing revealed hierarchical correspondence in mid-level feature processing, but not low- and high-level feature processing, further clarifying the gap between CNNs and human brains in visual scene processing.

## 5. Data and code availability

The neural data, stimuli and ground-truth annotations are uploaded on https://osf.io/7c9bz/. The code used for this project is under https://github.com/Agnessa14/Mid-level-features.

## 6. Acknowledgments

A.K. is supported by a Ph.D. fellowship provided by the European Research Council (ERC) starting grant (ERC-StG-2018-803370) and by a stipend from the Einstein Center for Neurosciences. R.M.C. is supported by German Research Council (DFG) grants (CI 241/3-1, CI 241/1-7) and the European Research Council (ERC) starting grant (ERC-StG-2018-803370). G.R. is supported by the DFG Research Unit grant (FOR 5368). We thank the HPC Service of ZEDAT, Freie Universität Berlin (Bennett et al., 2020) for computing time. We thank Antoniya Boyanova and Mirjam Marx for helping with data acquisition and Alessandro Gifford, Johannes Singer, Daniel Janini and Siying Xie for their valuable comments on the manuscript.

## 7. Author contributions

A.K., A.L., V. B., K.D., G.R. and R.M.C conceived and designed the experiment and the analyses. M.P. created the stimulus set. A.K., A.L. and V.B. performed the experiments. R.L. contributed analysis code. A.K., A.L. and V.B. analysed the data. R.M.C. and K.D. supervised the analysis. A.K., A.L. and R.M.C. wrote the manuscript. R.M.C., K.D. and G.R. reviewed and edited the manuscript.

