## Supplementary file for "Investigating the temporal dynamics and modelling of mid-level feature representations in humans"

**Supplementary material**

**Supplementary Table 1. Number of principal components used to capture 90% or more of the variance in the CNN encoding analysis.**

| | **Number of components** |
|---|---|
| **Image CNN** | |
| Layer 1.0 | 778 |
| Layer 1.1 | 837 |
| Layer 2.0 | 844 |
| Layer 2.1 | 852 |
| Layer 3.0 | 763 |
| Layer 3.1 | 782 |
| Layer 4.0 | 789 |
| Layer 4.1 | 569 |
| **Video CNN** | |
| Layer 1.0 | 849 |
| Layer 1.1 | 864 |
| Layer 2.0 | 843 |
| Layer 2.1 | 846 |
| Layer 3.0 | 772 |
| Layer 3.1 | 782 |
| Layer 4.0 | 765 |
| Layer 4.1 | 625 |

**Supplementary Table 2. Statistical details for the encoding analysis on EEG data.**

| | **Significant time points (ms)** | **Peak latency [95% CI] (ms)** |
|---|---|---|
| **Images** | | |
| Edges | -80, 60:860 | 80 [80, 100] |
| Reflectance | 60:520 | 100 [100, 120] |
| Lighting | -60, 60:960 | 120 [120, 240] |
| World normals | 60:660 | 180 [140, 220] |
| Scene depth | 60:760 | 220 [120, 220] |
| Skeleton position | 60:980 | 260 [260, 520] |
| Action identity | -200, 80:980 | 460 [280, 540] |
| **Videos** | | |
| Edges | 60:860 | 100 [80, 100] |
| Reflectance | -260:-240, 60:680 | 100 [100, 100] |
| Lighting | 60:680, 740 | 200 [120, 220] |
| World normals | 60:700, 780 | 200 [80, 200] |
| Scene depth | -320, 60:760, 800 | 200 [120, 220] |
| Skeleton position | 60:980 | 220 [220, 220] |
| Action identity | -320, -120, 60:940, 980 | 280 [240, 440] |
| **Difference** | | |
| Edges | - | 20 [0, 20] |
| Reflectance | - | 0 [0, 20] |
| Lighting | - | 80 [0, 100] |
| World normals | - | 20 [0, 100] |
| Scene depth | - | 20 [0, 100] |
| Skeleton position | - | 40 [40, 300] |
| Action identity | - | 180 [20, 300] |

Significant time points, peak latencies with their 95% confidence intervals (10,000 bootstrap iterations over participants) and peak values for the encoding analysis on image and video EEG data.

**Supplementary Table 3. Statistical details for the encoding analysis on DNN activations.**

| | Significant layers | Peak latency [95% CI] |
|---|---|---|
| **Image CNN** | | |
| Edges | 1.1, 1.0, 2.0, 2.1, 3.0, 3.1, 4.0, 4.1 | 3.0 [1.0, 4.1] |
| Reflectance | 1.1, 1.0, 2.0, 2.1, 3.0, 3.1, 4.0, 4.1 | 1.0 [1.0, 3.0] |
| Lighting | 1.1, 1.0, 2.0, 2.1, 3.0, 3.1, 4.0, 4.1 | 1.0 [1.0, 4.1] |
| World normals | 1.1, 1.0, 2.0, 2.1, 3.0, 3.1, 4.0, 4.1 | 3.0 [3.0, 4.1] |
| Scene depth | 1.1, 1.0, 2.0, 2.1, 3.0, 3.1, 4.0, 4.1 | 3.1 [3.0, 4.1] |
| Skeleton position | 1.1, 1.0, 2.0, 2.1, 3.0, 3.1, 4.0, 4.1 | 4.1 [3.0, 4.1] |
| Action identity | 1.1, 1.0, 2.0, 2.1, 3.0, 3.1, 4.0, 4.1 | 3.1 [1.0, 4.1] |
| **Video CNN** | | |
| Edges | 1.1, 1.0, 2.0, 2.1, 3.0, 3.1, 4.0, 4.1 | 3.0 [2.0, 4.1] |
| Reflectance | 1.1, 1.0, 2.0, 2.1, 3.0, 3.1, 4.0, 4.1 | 1.0 [1.0, 4.1] |
| Lighting | 1.1, 1.0, 2.0, 2.1, 3.0, 3.1, 4.0, 4.1 | 3.0 [1.0, 3.1] |
| World normals | 1.1, 1.0, 2.0, 2.1, 3.0, 3.1, 4.0, 4.1 | 3.0 [3.0, 4.1] |
| Scene depth | 1.1, 1.0, 2.0, 2.1, 3.0, 3.1, 4.0, 4.1 | 3.1 [3.0, 4.0] |
| Skeleton position | 1.1, 1.0, 2.0, 2.1, 3.0, 3.1, 4.0, 4.1 | 4.1 [3.0, 4.1] |
| Action identity | 1.1, 1.0, 2.0, 2.1, 3.0, 3.1, 4.0, 4.1 | 4.1 [3.0, 4.1] |
| **Difference** | | |
| Edges | - | 0 [0, 5] |

| | | |
|---|---|---|
| Reflectance | 2.1 | 0 [0, 7] |
| Lighting | - | 4 [0, 5] |
| World normals | - | 0 [0, 3] |
| Scene depth | - | 0 [0, 3] |
| Skeleton position | - | 0 [0, 3] |
| Action identity | 2.1, 3.0, 3.1, 4.0, 4.1 | 2 [0, 7] |

Significant layers, peak latencies with their 95% confidence intervals and peak values for the encoding analysis on scene and action CNN activations. The confidence intervals were calculated by bootstrapping the results for the reduced unit activations (N=10 000).
